# Lemon: a framework for rapidly mining structural information from the Protein Data Bank

**DOI:** 10.1101/379891

**Authors:** Jonathan Fine, Gaurav Chopra

## Abstract

**Motivation:** The protein data bank (PDB) currently holds over 140,000 biomolecular structures and continues to release new structures on a weekly basis. The PDB is an essential resource to the structural bioinformatics community to develop software that mine, use, categorize, and analyze such data. New computational biology methods are evaluated using custom benchmarking sets derived as subsets of 3D experimentally determined structures and structural features from the PDB. Currently, such benchmarking features are manually curated with custom scripts in a non-standardized manner that results in slow distribution and updates with new experimental structures. Finally, there is a scarcity of standardized tools to rapidly query 3D descriptors of the entire PDB.

**Approach:** Our solution is the Lemon framework, a C++11 library with Python bindings, which provides a consistent workflow methodology for selecting biomolecular interactions based on user criterion and computing desired 3D structural features. This framework can parse and characterize the entire PDB in less than ten minutes on modern, multithreaded hardware. The speed in parsing is obtained by using the recently developed MacroMolecule Transmission Format (MMTF) to reduce the computational cost of reading text-based PDB files. The use of C++ lambda functions and Python binds provide extensive flexibility for analysis and categorization of the PDB by allowing the user to write custom functions to suite their objective. We think Lemon will become a one-stop-shop to quickly mine the entire PDB to generate desired structural biology features. The Lemon software is available as a C++ header library along with example functions at https://github.com/chopralab/lemon.

## 1 Introduction

Experimental structures deposited in the Protein Data Bank (PDB) (Rose *et al*., 2015) has resulted in several advances for structural and computational biology scientific and education communities. Several software packages have been developed using and applying data available in the PDB. Computational structural biology methods are evaluated using several benchmarking datasets mined from the PDB. As one example, for protein-ligand docking, the Astex (Hartshorn *et al*., 2007), PDBbind (Liu *et al*., 2017), and DUD-E (Mysinger *et al*., 2012) sets have been used to predict the 3D coordinates of ligands, rank target activity, and discriminate binders from non-binders.

Additionally, the knowledge-based forcefields for protein structure refinement (Chopra *et al*., 2008) and scoring functions used to evaluate ligand poses in a protein binding site (Bernard and Samudrala, 2009) require extensive feature mining of the PDB. The process for developing these benchmarking sets, structural features for knowledge-based forcefields and scoring functions are non-standard, time-consuming and computationally challenging as it requires significant computational resources to mine different 3D descriptors in the PDB. Development of software for mining these 3D features and use them for machine learning methods is challenging due to the increase in individual entry size as a significant computational cost is needed to parse large text-based formats.

The Macro Molecular Transmission Format (MMTF) (Bradley *et al*., 2017) was recently introduced to significantly reduce the time required to parse text-based formats traditionally used to store crystallographic data. MMTF requires a fraction of the computation time to read multiple files into computer memory as it uses an encoding format tailored specifically to protein and nucleic acid coordinate data and topology. Specifically, MMTF stores connectivity and chemical grouping data not captured in the PDB and mmCIF formats that are leveraged by Lemon’s data extraction framework. Lemon uses the entire PDB as Hadoop sequence files that are packaged as 578 independent subsets for all MMTF entries and used for the development of highly parallel workflows (Figure 1). Lemon is the only C++11 software package to our knowledge to parse the Hadoop sequence files natively.

**Fig. 1.**
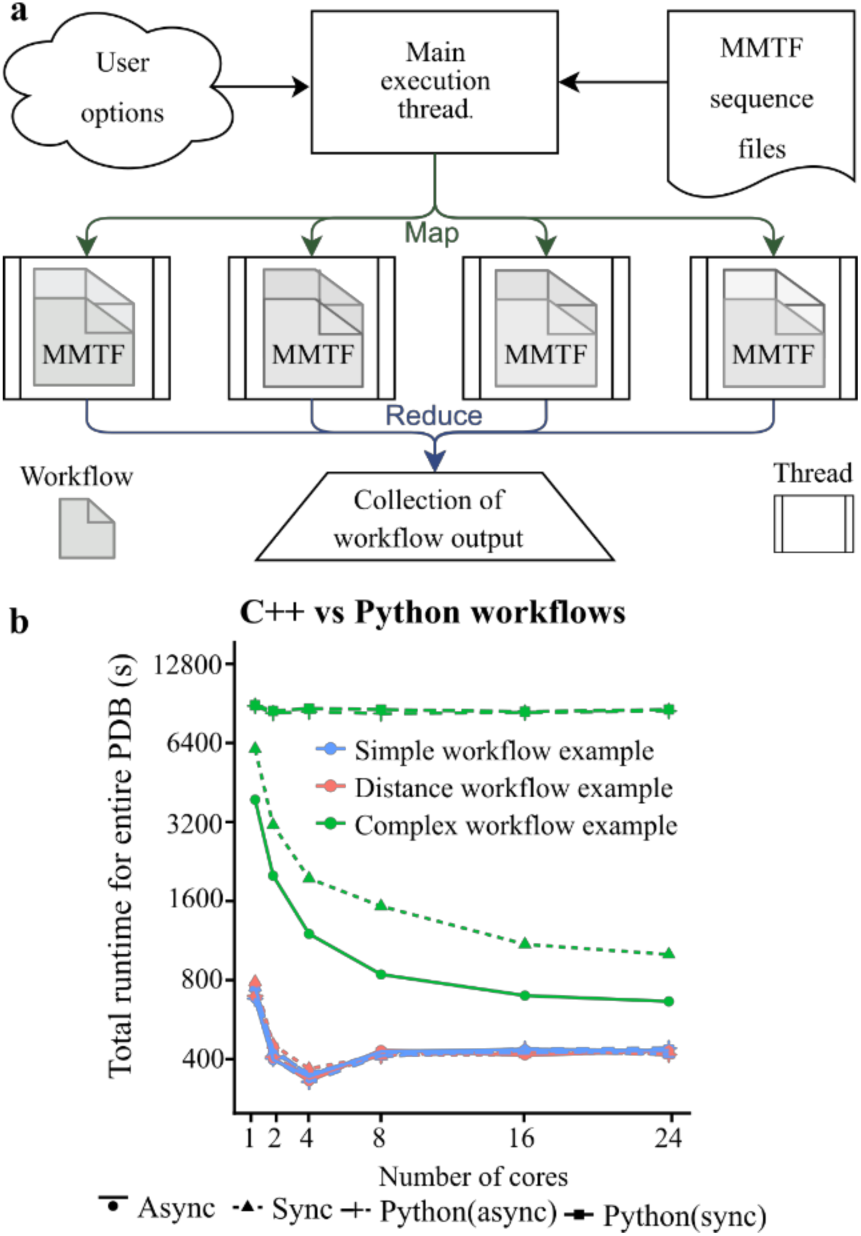
Workflow for Lemon. **a.** The overall work follow for the Lemon framework is given. The user provides C++ or Python API Lambda functions which use predefined functions to query information about each complex to filter the PDB into a desired subset. **b** A comparison between the C++ and Python benchmarking sets, showing the effect of multiple cores on overall runtime for simple to complex workflows for GCC (asynchronous, ‘Async’ and traditional or synchronous, ‘Sync’ threading).

### 2 Materials and methods

The Lemon framework uses a paradigm similar to MapReduce developed by Google for mining ‘Big Data’(Dean and Ghemawat, 2004). The user provides a ‘worker’ function that accepts two arguments: an object that represents the structure(s) of the chemical entities, and a string representing the four-letter PDBID. Lemon evaluates this function for all macromolecule entries in a multithreaded manner (Figure 1a), allowing one to perform any calculation on the structural information encoded by the MMTF file.

The MMTF object given to the user contains biomolecular data at the atomic, chemical group, and molecular levels. This includes the position, name, element type, and charge of the biomolecular atoms as well as the name, chain, biologic assembly, chemical links and composition type of chemical groups (e.g., protein residues). These features are examples that can be used to create workflows to select and extract desired 3D interactions.

Since a primary goal of the Lemon framework is to create standardized workflows, we have represented an example workflow pictorially (**Figure S1**). A workflow calculation is performed on the entire PDB database that is stored in its entirety on the user’s local machine. However, users can also choose to prefilter the database using a query generated on the RCSB website (see section **Using an RCSB search** in supporting information).

The workflow examples (Listings) are divided into ‘simple,’ ‘distance-based,’ and ‘complex’ categories based on the computational complexity of the workflow. First, the user ‘selects’ chemical groups present in the PDB entry using functions in Lemon for selecting small-molecules, metals, nucleic acids, amino acids, etc. These functions work on the group level by querying the group’s size and composition type. Additionally, it can also include the selection of topological information. Examples for these selectors are given in **Listing S1-S6**.

After obtaining a list of groups, the user can further divide (‘prune’) these groups based on 3D environment, biologic relevance, or frequency in the PDB. Lemon provides functions to find biologically identical groups, common groups (see **Table S1-S2**), and interacting groups via spatial relationship in 3D. Example lambda functions for ‘pruning’ groups are given in **Listings S7-S12**.

Finally, a workflow will calculate a feature of interest. For example, a user may perform structural alignment to a reference protein (**Listing S13**), calculate a docking score (**Listing S14**), or output statistics on geometries of bonded entities (**Listing S15-S18**). To show case the Python version of Lemon, three example workflows were ported to Python (**Listing S19-S21**). The information obtained from these workflows can then be directly used in machine learning approaches and the development of new structural biology methods.

Lemon also implements two different threading models based on the specifications of the C++ standard library. The first is a traditional (synchronous, ‘sync) threading approach which divides the PDB into 578 subsets and launches a user-defined number of threads to handle an equal portion of these 578 subsets (e.g., if the user selects two threads each thread will handle 289 subsets). The second is an asynchronous (‘async’) model that schedules 578 threads and executes a given number of them in parallel. Specifically, for async, the next queued thread executes when a thread completes, compared to the ‘sync’ model that requires all threads to complete.

## 3 Results and discussion

### 3.1 Querying the Protein Data Bank takes minutes

To measure Lemon’s execution time, we ran all example listings provided in the supporting information for different levels of multithreading and compiler architectures. The calculations were performed on a community cluster with each node consisting of two 12-core Intel Xeon Gold “Sky Lake” processors (see **Benchmarking Lemon** in supporting information). There are differences in computational time for a ‘simple,’ ‘distance-based,’ and ‘complex’ workflow (**Listings S6, S10, S18**) including the time to decompress and parse the MMTF files (**Figure S2**). The average runtime for all workflows with ‘async’ threading on eight cores (commodity hardware) takes ∼8 minutes to complete. The Lemon outputs for these queries are shown in **Figures S6-S8** and **Tables S3-S4.**

### 3.2 Workflow runtime influences threading efficiency

Asynchronous threading is more efficient for ‘complex’ workflows compared to sync threading (Figure 1b). Theoretically, the sync threading time should be more than async because it needs to wait for other threads to complete. However, the async and sync runtimes are similar for ‘simple’ and ‘distance-based’ workflows (**Listings S6, S10**) but differ for complex workflow (**Listing S18**) for 2 and 4 cores (**Figure S3**). The runtime reduces with increase in the number of cores (see 1, 2, and four cores in **Figure S4**). However, for some simple and distance-based workflows runtime increased from 4 cores to 8 cores (**Figure S4**). This result may be due to increased performance penalty for atomic (thread locking) operations after completion of each thread. This *hypothesis* is supported by the continued increase in performance for ‘complex’ operations as they are less likely to become bound.

### 3.3 Large biomolecules do not affect runtime

**Figure S4** shows that removal of the largest size PDBs (3J3Q, 3J3Y, 5Y6P) does not significantly reduce the overall runtime for most workflows when compared to the entire PDB (left column in the figure). An exception is the calculation of small-molecule/peptide interactions that requires distance calculations between millions of atoms for large complexes (see Peptides in **Figure S4**). Hence, Lemon workflows scale with the size PDB entries. This is a significant result given the increase in the amount of large structures in the PDB (RCSB stats page).

### 3.4 Compiler choice significantly impacts runtime

The selection of the C++ compiler dramatically affects the performance of Lemon (**Figure S5**). However, the timings shown in **Figure S5** indicate that there is only a marginal difference between the ‘sync’ and ‘async’ models averaged over all workflows. The GNU Compiler Collection (GCC) version 6.3.0 with ‘sync’ threading compilation outperforms the Intel compiler version 17.0.1.132 with sync threading (**Figure S5**, green and blue bars). This discrepancy could be a result of GCC’s use of a modern version of the C++ standard library or the specific optimizations performed by this compiler are better for Lemon. Further profiling is beyond the scope of this work and may be addressed in future publications.

### 3.5 Python is slower than C++ for complex workflows

The data shown in Figure 1b indicates that the Python bindings are just as fast as the C++ version for ‘simple’ and ‘distance-based’ workflows. Complex calculations scale poorly with the number of cores, a result due to the Python global interpreter lock. This underlines the importance of development in the C++ language, potentially after prototyping a complex workflow in Python.

### 3.6 Code availability

Lemon is hosted on GitHub (see ‘**Obtaining Lemon**’ in supporting information) along with C++ and Python API documentation on the GitHub page repository. File input and output are provided by the Chemfiles library. A link to the Lemon GitHub repository has been added to the official MMTF webpage on mmtf.rcsb.org.

## Acknowledgements

We thank Gerardo Tauriello and Daniel Farrell for MMTF-CPP and Guillaume Fraux for Chemfiles libraries.

