## Supporting Information for "Lemon: a framework for rapidly mining structural information from the Protein Data Bank"

#### Obtaining Lemon

The Python version of Lemon is available on the PyPI package repository and can be installed on Windows, MacOS and Linux using the following command. Note that this only installs the Python version of Lemon and does provide access to the C++ API.

```
$ python3 -m pip install --user candiy-lemon
```

The lemon benchmarking framework is written in C++11. Its only dependencies are a C++11 compiler, the CMake tools, and the Chemfiles library.  Note that the Chemfiles library will be automatically installed by the lemon install scripts if it is not already installed your system. Note that Lemon itself is a 'header-only' library, but users will also need a copy of the Chemfiles library to include it in their projects.

There are two threading methodologies that can be used by Lemon.  The default is based on *std::thread* provided by the C++11 standard and should compile with any C++11 compiler.  A second methodology uses a thread pool which uses *std::async* and requires C++14 features.

Lemon has been freely released on GitHub. To obtain the software, complete the following steps:

```
$ git clone https://github.com/chopralab/lemon.git
$ cd lemon/
$ mkdir build/
$ cd build
$ cmake .. -DCMAKE_BUILD_TYPE=Release
$ make
```

The example binaries will be created in the 'progs' subdirectory of `build`.  It is recommended that you supply the 'make' command with an additional argument '-j ##' where '##' is the number of physical cores on your machine.  To use the *std::async* version of Lemon, add `-DLEMON_TEST_ASYNC=ON` to the cmake configuration line.  To build documentation, add `-DLEMON_BUILD_DOCs=ON` to the configuration line.  An online version of this documentation is available at https://chopralab.github.io/lemon/latest/.

### Using an RCSB search

Lemon supports the ability to skip user selected entries using criteria created using the advanced search functionality on the RCSB website. This procedure is detailed below:

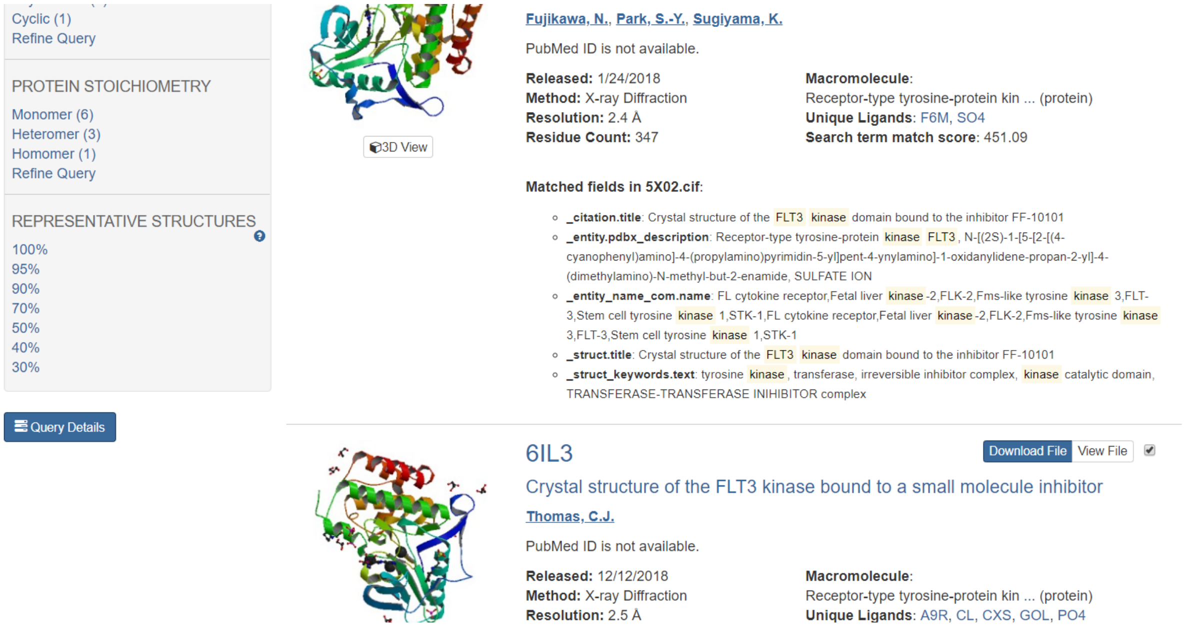


First, perform a search on the RCSB website and click the query details button in the bottom left hand corner of the search results page.

**Query in XML format**


```
<orgPdbQuery>
  <version>head</version>
  <queryType>org.pdb.query.simple.AdvancedKeywordQuery</queryType>
  <description>Text Search for: flt3 kinase</description>
  <queryId>6FBC1D79</queryId>
  <resultCount>10</resultCount>
  <runtimeStart>2019-02-02T19:53:03Z</runtimeStart>
  <runtimeMilliseconds>208</runtimeMilliseconds>
  <keywords>FLT3 Kinase</keywords>
</orgPdbQuery>
```


Copy and paste the XML Query in to a text file and run the script *obtain_entries_from_search.pl* (included in the scripts directory) to produce an entries file (extension does not matter). You can provide this file to Lemon workflows using the '-e' and '-s' command line arguments. The former will launch a lemon workflow that only includes the selected entries and the latter will skip the selected entries when performing a workflow. These options can be used together.

#### Benchmarking Lemon

All Lemon benchmarks were run on the Brown high performance computing cluster. The nodes that comprise this cluster have 96 Gb of memory and two Sky Lake CPUs clocked at 2.60GHz, resulting in a total of 24 cores. Further details about this compute cluster can be found at https://www.rcac.purdue.edu/compute/brown.

The calculation of the timings for individual workflows versus the overall runtime was done by using the LEMON_BENCHMARK flag during compilation. These timings include decompression of the MMTF file using the Gzip algorithm but include neither the time required to read the compressed MMTF from the Hadoop sequence file into memory, nor the time required to output the results. These timings are printed to STDERR after the workflow completes for a single entry.

This procedure was performed three times using different compilation settings to understand the difference between these settings and overall workflow runtime. The three settings are (1) using the Intel C++ Compiler 17.0.1.132 with traditional (synchronous) threading enabled, (2) GNU C++ Compiler 6.3.0 with traditional threading, and (3) GNU C++ Compiler 6.3.0 with asynchronous (async) threading enabled.

Lemon jobs were submitted to the cluster's using the following script when different processor counts where supplied during submission. All benchmarking was performed on a single node.

```
#!/usr/bin/env bash
#PBS -d .
#PBS -l walltime=04:00:00

if [[ -z $LEMON_PROG ]]
then
    echo "You must specify the LEMON_PROG variable"
    exit 1
fi

PPN=$(wc -l $PBS_NODEFILE | cut -f1 -d’ ‘)

# /dev/shm is the location of shared memory on RedHat systems.
# you may need to change this location!
tar -xf full.tar -C /dev/shm/

SECONDS=0
```

```
time $RCAC_SCRATCH/lemon/build_${compiler}/bin/lemon/$LEMON_PROG -w /dev/shm/full
-n $PPN > ${LEMON_PROG}.log

echo "$LEMON_PROG $PPN $compiler $SECONDS"


rm -fr /dev/shm/full
```

**Table S1:** Common chemical groups ignored in the above examples.

| Three letter code | Full name |
|---|---|
| **CIT** | Citric Acid |
| **FLC** | Citric Acid Ion |
| **ICT** | Isocitrate |
| **SAM** | S-Adenosylmethionine |
| **SIA** | O-Sialic Acid |
| **CLA** | Chlorophyll |
| **HEM** | PROTOPORPHYRIN IX CONTAINING FE |
| **HEA** | Heme-A |
| **HEB** | Heme-B/C |
| **HEC** | Heme-C |
| **MES** | 2-(N-MORPHOLINO)-ETHANESULFONIC ACID |
| **EPE** | 4-(2-HYDROXYETHYL)-1-PIPERAZINE ETHANESULFONIC ACID |
| **GOL** | Glycerol |
| **FAD** | FLAVIN-ADENINE DINUCLEOTIDE |
| **FMN** | FLAVIN MONONUCLEOTIDE |
| **NAD** | NICOTINAMIDE-ADENINE-DINUCLEOTIDE |
| **NAP** | NADP NICOTINAMIDE-ADENINE-DINUCLEOTIDE PHOSPHATE |
| **ADP** | ADENOSINE-5'-DIPHOSPHATE |
| **ATP** | ADENOSINE-5'-TRIPHOSPHATE |
| **GDP** | GUANOSINE-5'-DIPHOSPHATE |
| **GTP** | GUANOSINE-5'-TRIPHOSPHATE |
| **DTP** | 2'-DEOXYADENOSINE 5'-TRIPHOSPHATE |
| **BE7** | (4-CARBOXYPHENYL)(CHLORO)MERCURY |
| **MHA** | (CARBAMOYLMETHYL-CARBOXYMETHYL-AMINO)-ACETIC ACID |
| **DHD** | 2,4-DIOXO-PENTANEDIOIC ACID |
| **B3P** | 2-[3-(2-HYDROXY-1,1-DIHYDROXYMETHYL-ETHYLAMINO)-PROPYLAMINO]-2-HYDROXYMETHYL-PROPANE-1,3-DIOL |
| **BTB** | 2-[BIS-(2-HYDROXY-ETHYL)-AMINO]-2-HYDROXYMETHYL-PROPANE-1,3-DIOL |
| **NHE** | 2-[N-CYCLOHEXYLAMINO]ETHANE SULFONIC ACID |

**Table S2:** Fatty acids and other common molecules present throughout the PDB.

| Three letter code | Full name |
|---|---|
| **PG4** | TETRAETHYLENE GLYCOL |
| **PG5** | 1-METHOXY-2-[2-(2-METHOXY-ETHOXY]-ETHANE |
| **PG6** | 1-(2-METHOXY-ETHOXY)-2-{2-[2-(2-METHOXY-ETHOXY]-ETHOXY}-ETHANE |
| **1PE** | PENTAETHYLENE GLYCOL |
| **2PE** | NONAETHYLENE GLYCOL |
| **1PG** | 2-(2-{2-[2-(2-METHOXY-ETHOXY)-ETHOXY]-ETHOXY}-ETHOXY)-ETHANOL |
| **PE4** | 2-{2-[2-(2-{2-[2-(2-ETHOXY-ETHOXY)-ETHOXY]-ETHOXY}-ETHOXY)-ETHOXY]-ETHOXY}-ETHANOL |
| **PE8** | 3,6,9,12,15,18,21-HEPTAOXATRICOSANE-1,23-DIOL |
| **PG6** | 1-(2-METHOXY-ETHOXY)-2-{2-[2-(2-METHOXY-ETHOXY]-ETHOXY}-ETHANE |
| **P33** | 3,6,9,12,15,18-HEXAOXAICOSANE-1,20-DIOL |
| **C8E** | (HYDROXYETHYLOXY)TRI(ETHYLOXY)OCTANE |
| **XPE** | 3,6,9,12,15,18,21,24,27-NONAOXANONACOSANE-1,29-DIOL |
| **N8E** | 3,6,9,12,15-PENTAOXATRICOSAN-1-OL |
| **DR6** | 3,6,9,12,15-PENTAOXATRICOSAN-1-OL |
| **MYR** | MYRISTIC ACID |
| **OTE** | 2-{2-[2-(2-OCTYLOXY-ETHOXY)-ETHOXYL]-ETHOXY}ETHANOL |
| **OLA** | OLEIC ACID |
| **OLB** | (2S)-2,3-dihydroxypropyl (9Z)-octadec-9-enoate |
| **OLC** | (2R)-2,3-dihydroxypropyl (9Z)-octadec-9-enoate |
| **PLM** | PALMITIC ACID |
| **PAM** | PALMITOLEIC ACID |
| **PEE** | 1,2-Dioleoyl-sn-glycero-3-phosphoethanolamine |
| **LHG** | 1,2-DIPALMITOYL-PHOSPHATIDYL-GLYCEROLE |
| **MC3** | 1,2-DIMYRISTOYL-RAC-GLYCERO-3-PHOSPHOCHOLINE |
| **SPM** | SPERMINE |
| **SPK** | SPERMINE (FULLY PROTONATED FORM) |
| **SPD** | SPERMIDINE |

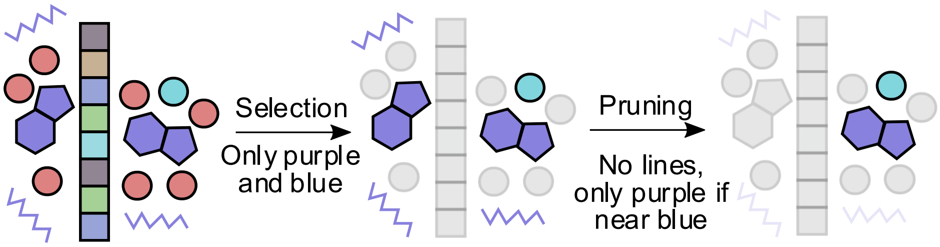


**Figure S1**: Diagram showing the recommended Lemon workflow. The workflow begins with selection when the user provides criterion on which chemical groups, they wish to perform calculations on. In this example, the purple groups represent small molecules, the red groups represent water, the cyan groups represent metals, and the boxed groups represent amino acids. Here, the user has selected small-molecules and metal ions. The next step is pruning of the selected residues. Here, the user has decided to remove the small molecules which do not contain rings and remove small molecules which are not within proximity of a metal ion. Finally, the user can perform a calculation on their selected pairs.

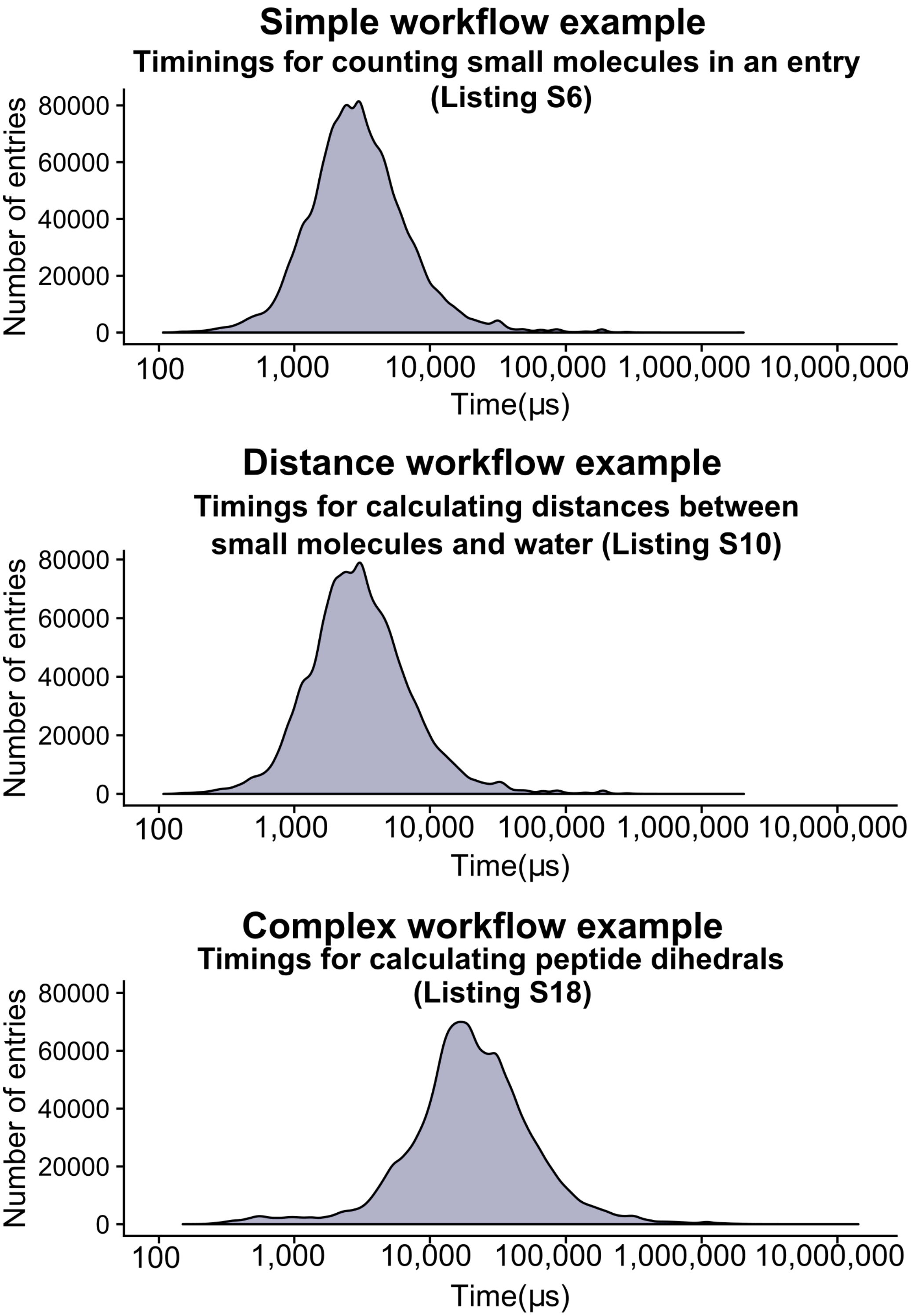


**Figure S2:** Timings for individual workflows are given as examples from the three different types of workflows. These times were taken from a single core launch to ensure that each timing was as independent of other calculations. These results indicate there is little difference between the 'simple' and 'distance-based' calculations, a potential result of the reduced computational cost due to carefully 'selecting' and 'pruning' chemical groups before performing the distance calculation.

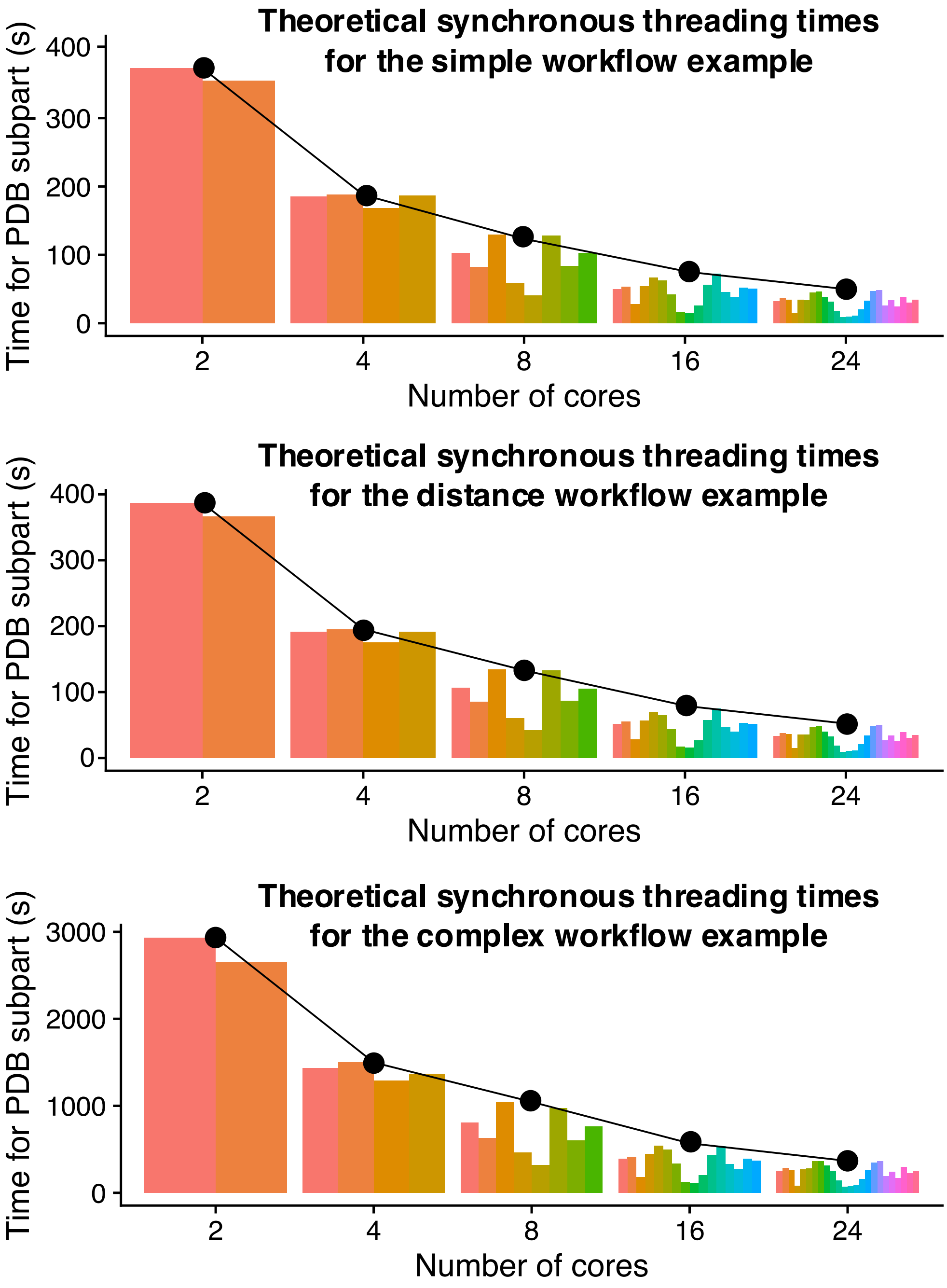


**Figure S3:** Theoretical minimum performance of each traditional thread as computed for the three examples shown in **Figure S2**. These are calculated by grouping the individual entries by their subgroup and summing the total time. The result is the colored subpart, which is dependent on the number of cores executed by the user as this number is used to calculate the total number of subparts. This plot shows that the maximum runtime of a subpart decreases as the number of cores increase for all three example operations (black line). It also indicates that the time taken by each sub part is the same for 2 and 4 cores but diverges for higher core counts.

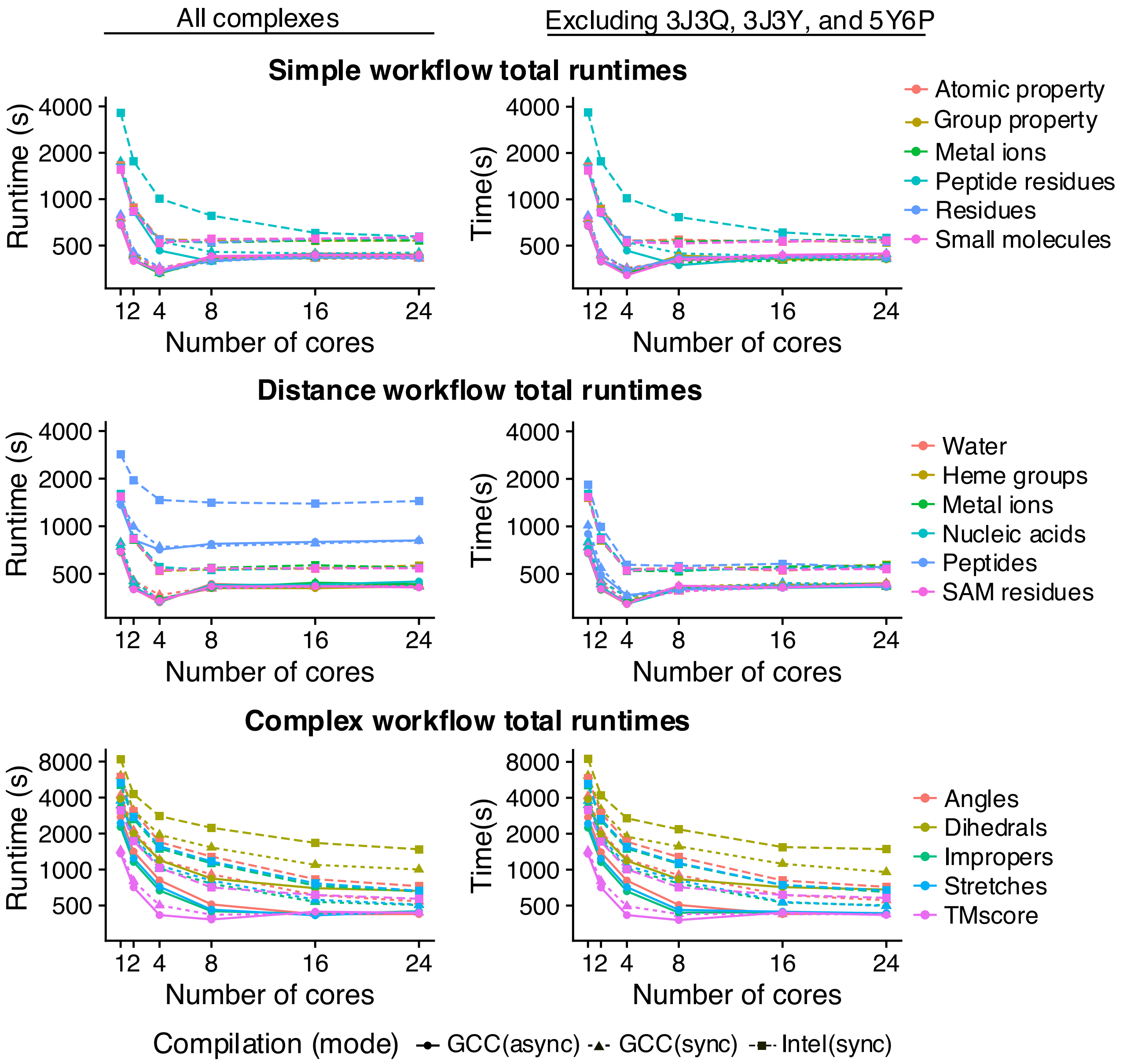


**Figure S4:** Benchmarking results for the Lemon workflows listed previously in this document. Here, we have divided these workflows by their relative complexity. We ran the benchmarking set for the entire PDB with (left column) and without the three largest size PDB entries, 3J3Q, 3J3Y, and 5Y6P (right column). These entries have a processing time at least 3 times greater than the remaining entries. Note that runtimes given in the Y-axis are plotted logarithmically. These plots show that 4 cores provide the optimal run time for 'simple' and 'distance-based' operations. Additional cores do improve runtime for 'complex' operations, however, indicating the possibility of an Input-Output bottleneck on fast calculations.

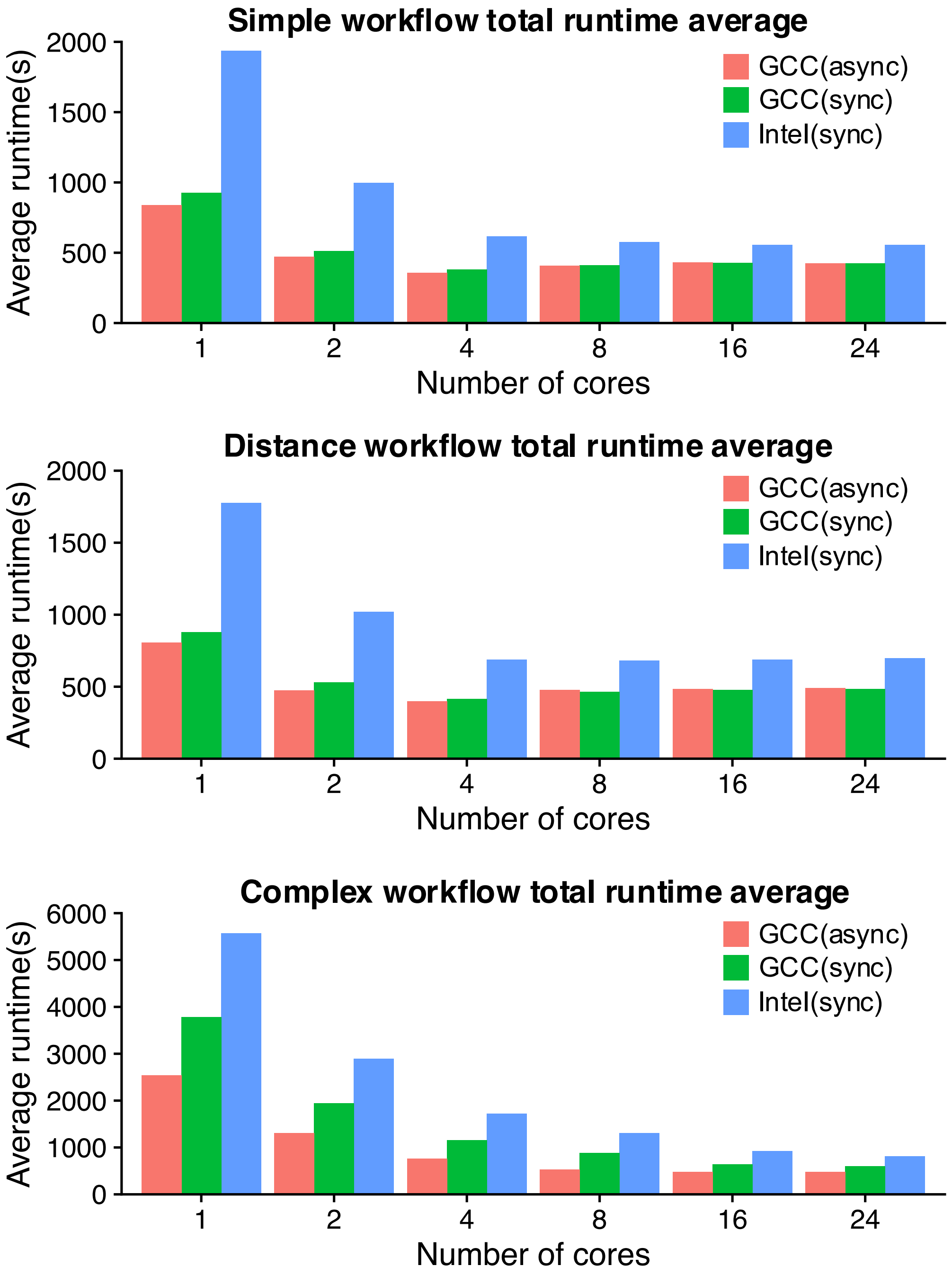


**Figure S5:** The average runtimes for the three different compiled versions of Lemon. These data show that overall the GCC compiler out performed the Intel compiler int all test cases. Further, they show the ‘async’ threading model is only marginally faster for the ‘simple’ and ‘distance-based’ workflows but holds improvements for the ‘complex’ calculations.

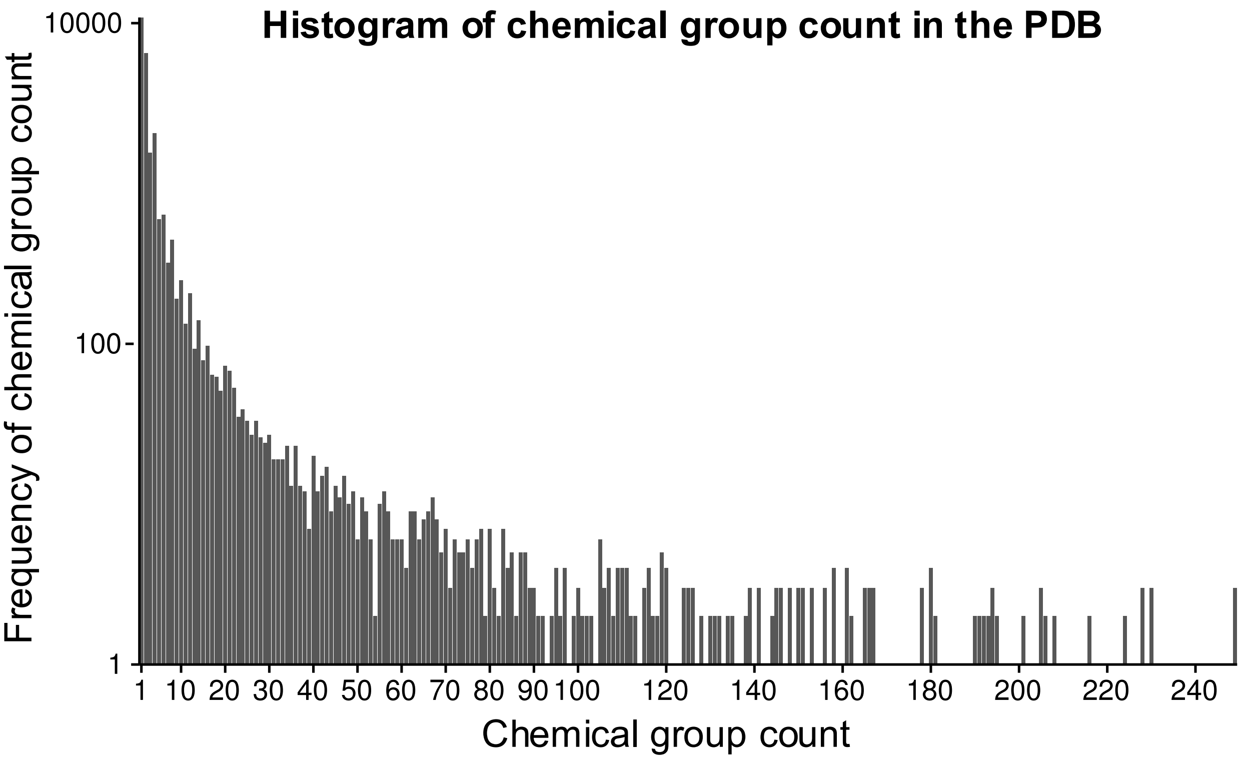


**Figure S6:** Histogram showing the frequency of a given chemical group count (maximum of 250). The X-axis is the chemical group count. This count is independent of chemical environment is determined from the three-letter code given to chemical groups in the PDB. For example, if the residue 'CFF' occurs once in PDBID 142N and thrice in PDBID 1L59, and occurs nowhere else in the PDB, then it has a count of 4. The Y-axis gives the frequency for all chemical group counts in the PDB. From this data we can conclude that the majority of chemical groups occur only once in throughout the entire PDB.

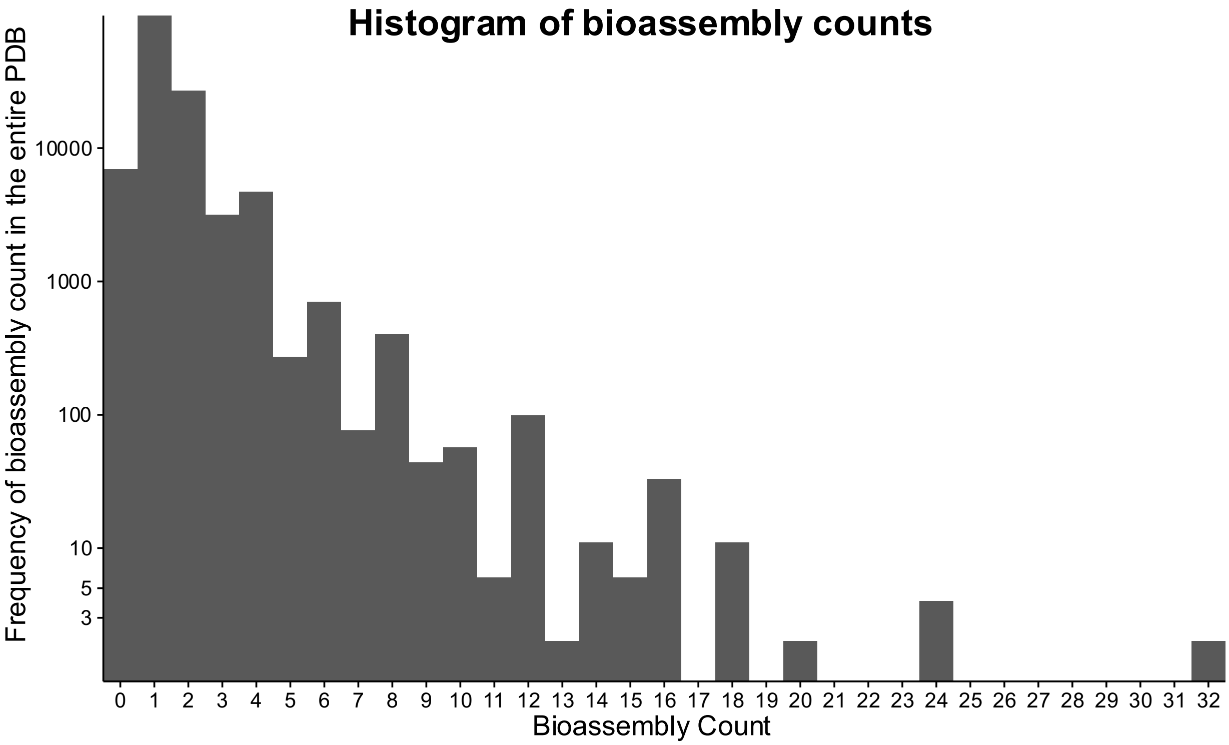


**Figure S7:** Histogram showing the frequency of bioassemblies (as defined by the depositor of a PDB file) throughout the PDB.

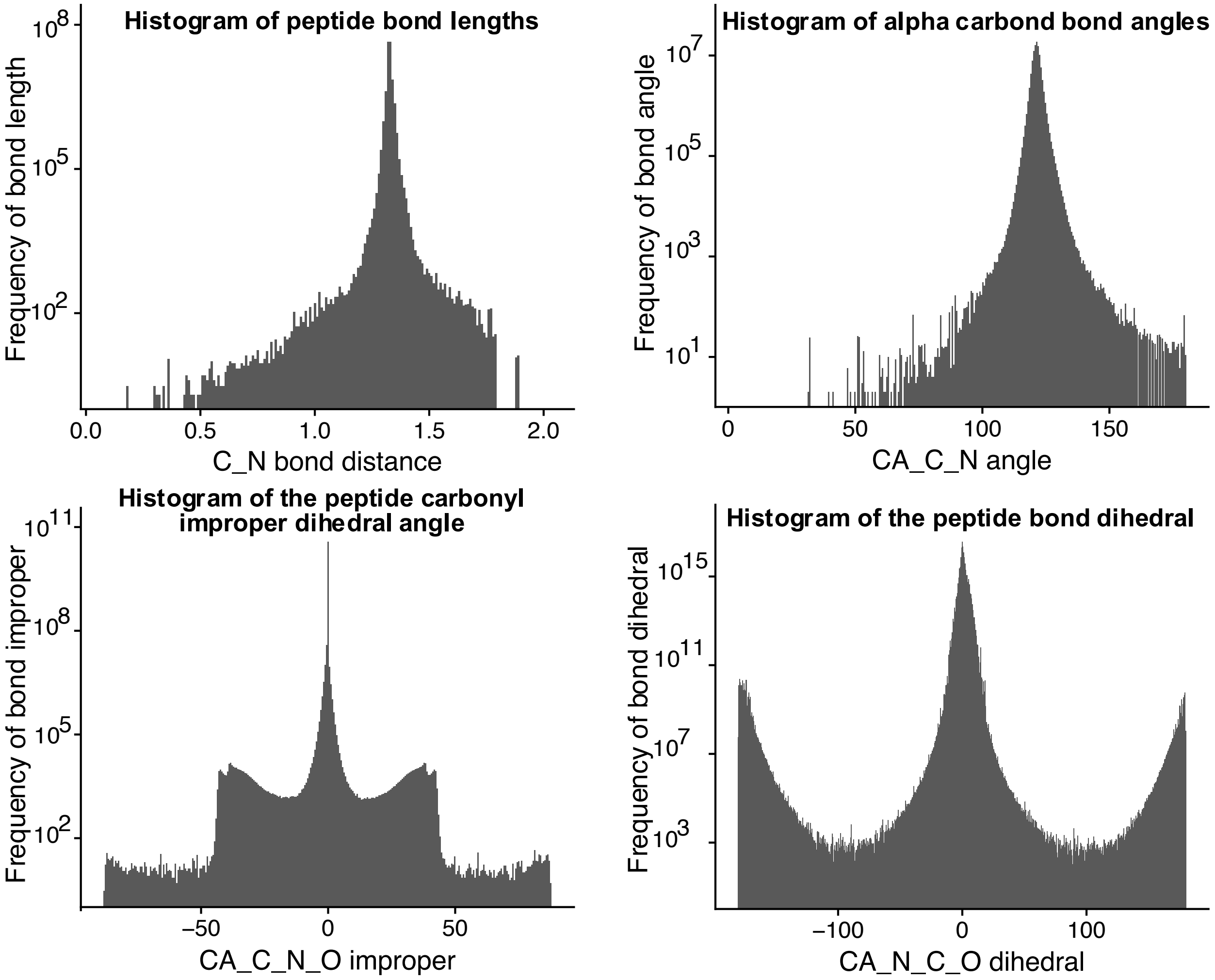


**Figure S8:** Histograms of various geometries centered around the peptide bond. These plots illustrate Lemon's ability to mine geometrical data from the PDB.

**Table S3:** Number of PDB entries which contain a given metal ion.

| Metal Element | Number of entries |
|---|---|
| **Li** | 72 |
| **Be** | 1 (4P4K) |
| **Na** | 7537 |
| **Mg** | 13418 |
| **K** | 2435 |
| **Ca** | 10197 |
| **V** | 1 (1QYL) |
| **Mn** | 3138 |
| **Fe** | 2175 |
| **Cu** | 1432 |
| **Hg** | 477 |

**Table S4:** Compounds which interact with SAM.

| PDBID | Three letter code | Count |
|---|---|---|
| **1P7L** | PPK | 2 |
| **5KJK** | 6T1 | 1 |
| **5KJM** | 6TM | 1 |
| **5KJN** | 6TL | 1 |
| **4E47** | 0N6 | 1 |
| **2G70** | HNT | 1 |
| **2G72** | F21 | 1 |
| **1R30** | DTB | 2 |
| **1RG9** | PPK | 4 |
| **2ZVJ** | KOM | 1 |
| **3A7E** | DNC | 1 |
| **5LSA** | DNC | 1 |
| **3PFG** | TLO | 1 |
| **1H1D** | BIA | 1 |
| **1VID** | DNC | 1 |
| **3BWM** | DNC | 1 |
| **3BWY** | DNC | 1 |
| **3BXO** | UPP | 2 |
| **3S68** | TCW | 1 |
| **3S7B** | NH5 | 1 |
| **4JDS** | 1L4 | 1 |
| **4JLG** | 1L8 | 1 |
| **3DMH** | GMP | 1 |
| **1XDS** | DRA | 2 |
| **4KIC** | PPY | 2 |
| **2PXC** | G3A | 1 |
| **3EMB** | GTG | 1 |
| **4M7T** | 25W | 1 |
| **2BR4** | P4C | 3 |
| **2CL5** | BIE | 1 |
| **4NDN** | PPK | 2 |
| **3I5U** | 5NA | 2 |
| **4NJG** | HHS | 2 |
| **4NJH** | 2K8 | 2 |
| **4NJI** | 2K8 | 2 |
| **4NJJ** | 2K8 | 2 |
| **4NJK** | 2KA | 2 |
| **4A6E** | ASE | 1 |
| **4ODJ** | 3PO | 1 |

| | | |
|---|---|---|
| **5T8S** | 3PO | 1 |
| **4RVG** | TYD | 1 |
| **5UL4** | B12 | 1 |
| **5V37** | 8WD | 1 |
| **5V3H** | 8WG | 1 |
| **5W8A** | A1S | 1 |
| **5WBV** | 9ZY | 1 |
| **4X61** | 3XV | 1 |
| **4XUC** | 43G | 1 |
| **4XUD** | 43H | 1 |
| **4XUE** | 43J | 1 |
| **4YND** | 4GQ | 1 |
| **5A1I** | AND, PPK | 1, 1 |
| **5ARF** | I9H | 1 |
| **5ARG** | H41 | 1 |
| **5AYF** | C7H | 1 |
| **5CCL** | 4ZW | 1 |
| **5CCM** | 4ZX | 1 |
| **5CPR** | 539 | 1 |
| **5EML** | 5QK | 1 |
| **5FEP** | 41K | 1 |
| **5FES** | 9SE | 1 |
| **5FHQ** | DNC | 1 |
| **5FHR** | DNC | 2 |

#### Example Workflows in C++ (Listings)

The following Lemon workflows are designed to teach a user how to write and design a workflow based on the recommended design pattern outlined in **Figure S1**.  While their purpose is primarily didactic, they can be repurposed into workflows which extract meaningful data from the PDB in a systematic and organized fashion.

We have classified these 18 benchmarks as ‘simple’, ‘distance-based’ and ‘complex’, based on the type of calculation performed by the workflow. The ‘simple’ operations do not involve computing any 3D geometry functions and are designed to show the ‘selection’ stage of a Lemon workflow. The ‘distance-based’ workflows involve the calculation of various distances between selected small-molecules and other selected chemical groups throughout the PDB.  These operations showcase the distance-based ‘pruning’ operations available in the Lemon API. Finally, the ‘complex’ calculations involve calculating complex geometric features such as angles, dihedrals, and improper dihedrals. They also include the calculation of all protein bonds and structural alignment.

**Listing S1:** (simple) C++ Lambda function to count the number of biological assemblies in the PDB. This example illustrates how to obtain information about a residue/group property (in this case symmetry) which could be used to determine if the user wishes to continue calculation.

```
int main(int argc, char* argv[]) {
    lemon::Options o(argc, argv);
    auto worker = [](chemfiles::Frame entry, const std::string& pdbid) {
        // Desired info is obtained directly, no pruning
        auto result = lemon::count::residue_property(entry, "assembly");

        // Output phase
        return pdbid + " " + std::to_string(result) + "\n";
    };

    auto collector = lemon::print_combine(std::cout);
    return lemon::launch(o, worker, collector);
}
```

**Listing S2:** (simple) C++ Lambda function to determine the number of alternative atom locations in all PDB entries.  This example illustrates the use of an atomic property to potentially screen entries which do not contain alternative locations.

```
int main(int argc, char* argv[]) {
    lemon::Options o(argc, argv);
    auto worker = [](chemfiles::Frame entry, const std::string& pdbid) {
        // Desired info is obtained directly
        auto result = lemon::count::atom_property(entry, "altloc");

        // Output phase
        return pdbid + " " + std::to_string(result) + "\n";
    };

    auto collector = lemon::print_combine(std::cout);
    return lemon::launch(o, worker, collector);
}
```

**Listing S3:** (simple) C++ Lambda function to select metal ions in the PDB. This workflow shows how the selection phase of a Lemon workflow works by filling a generic STL container with the desired residue ids. The output is the pdbid followed by all the metal ions found in the corresponding entry.

```
int main(int argc, char* argv[]) {
    lemon::Options o(argc, argv);
    auto worker = [](chemfiles::Frame entry,
                     const std::string& pdbid) {

        // Selection phase
        std::list<size_t> metal_ids;
        lemon::select::metal_ions(entry, metal_ids);

        // No pruning, straight to output phase
        return pdbid + lemon::count::print_residue_names(entry, metal_ids);
    };

    auto collector = lemon::print_combine(std::cout);
    return lemon::launch(o, worker, collector);
}
```

**Listing S4:** (simple) C++ Lambda function to determine the occurrence of all residues in the PDB. The purpose of this workflow is to show that one can return more than strings from a C++ lambda function as long they use a different 'combine' function object to handle this return value. The concept of 'combine' functions is detailed in the online documentation along with this example to illustrate it. It outputs all three-letter residue names and the number of times each is found throughout all entries in the PDB. Note that residues may occur multiple times in a single entry and this is reflected in this lambda.

```
int main(int argc, char* argv[]) {
    lemon::Options o(argc, argv);

    auto worker = [](chemfiles::Frame entry, const std::string&) {
        // Desired info is calculated directly, no pruning, output is done later
        lemon::ResidueNameCount rnc;
        lemon::count::residues(entry, rnc);
        return rnc;
    };

    lemon::ResidueNameCount resn_total;
    auto collector = lemon::map_combine<lemon::ResidueNameCount>(resn_total);
    lemon::launch(o, worker, collector);

    for (auto i : resn_total) {
        std::cout << i.first << "\t" << i.second << "\n";
    }
}
```

**Listing S5:** (simple) This example workflow combines concepts from the past two workflows to show that 'selection' can be combined with other workflow concepts via the separate functionality. Separate allows one to create a subset of an entry and perform further calculations on just the subset. This workflow is similar to **Listing S4**, but only prints residues with peptide linkage.

```
int main(int argc, char* argv[]) {
    lemon::Options o(argc, argv);
    auto worker = [](chemfiles::Frame entry, const std::string&) {
        lemon::ResidueNameCount rnc;
        // Selection phase
        chemfiles::Frame protein_only;
        auto peptides = lemon::select::peptides(entry);

        if (peptides.size() == 0) {
            return rnc;
        }
        lemon::separate::residues(entry, peptides, protein_only);
        // Output phase
        lemon::count::residues(protein_only, rnc);
        return rnc;
    };
    lemon::ResidueNameCount resn_total;
    auto collector = lemon::map_combine<lemon::ResidueNameCount>(resn_total);
    lemon::launch(o, worker, collector);
    for (auto i : resn_total) {
        std::cout << i.first << "\t" << i.second << "\n";
    }
}
```

**Listing S6:** (simple) This workflow is designed to introduce pruning to the user. In this specific example, selected small molecules are pruned by removing common cofactors and common fatty acids. No detailed calculations are performed yet, but such calculations will be introduced in the next workflows.

```
int main(int argc, char* argv[]) {
    lemon::Options o(argc, argv);

    auto worker = [](chemfiles::Frame entry,
                     const std::string& pdbid) {

        // Selection phase
        std::list<size_t> smallm;
        if (lemon::select::small_molecules(entry, smallm) == 0) {
            return std::string("");
        }

        // Pruning phase
        lemon::prune::identical_residues(entry, smallm);
        lemon::prune::cofactors(entry, smallm, lemon::common_cofactors);
        lemon::prune::cofactors(entry, smallm, lemon::common_fatty_acids);

        // Output phase
        return pdbid + lemon::count::print_residue_names(entry, smallm);
    };

    auto collector = lemon::print_combine(std::cout);
    return lemon::launch(o, worker, collector);
}
```

**Listing S7:** (distance-based) C++ Lambda function to determine the number of small molecules which interact with a metal ion within a distance cutoff. This workflow is designed to show how to select two different groups and perform a distance-based pruning operation on the two groups. It also introduces the concept of obtaining command-line arguments.

```
int main(int argc, char* argv[]) {
    lemon::Options o;
    auto distance = 6.0;
    o.add_option("--distance,-d", distance,
                 "Largest distance between a metal and a small molecule.");
    o.parse_command_line(argc, argv);
    auto worker = [distance](chemfiles::Frame entry,
                             const std::string& pdbid) {
        // Selection phase
        auto metals = lemon::select::metal_ions(entry);
        auto smallm = lemon::select::small_molecules(entry);
        // Pruning phase
        lemon::prune::identical_residues(entry, smallm);
        lemon::prune::cofactors(entry, smallm, lemon::common_cofactors);
        lemon::prune::cofactors(entry, smallm, lemon::common_fatty_acids);
        lemon::prune::keep_interactions(entry, smallm, metals, distance);
        // Output phase
        return pdbid + lemon::count::print_residue_names(entry, smallm);
    };
    auto collector = lemon::print_combine(std::cout);
    return lemon::launch(o, worker, collector);
}
```

**Listing S8:** (distance-based) C++ Lambda function to determine the number of small molecules which interact with a Heme group within a distance cutoff. This is similar to the last workflow and illustrates how selectors can be used to find cofactors instead of metal ion.

```
int main(int argc, char* argv[]) {
    lemon::Options o;
    auto distance = 6.0;
    o.add_option("--distance,-d", distance,
                 "Largest distance between the Heme and a small molecule.");
    o.parse_command_line(argc, argv);
    auto worker = [distance](chemfiles::Frame entry,
                             const std::string& pdbid) {
        // Selection phase
        auto hemegs = lemon::select::specific_residues(
            entry, {"HEM", "HEA", "HEB", "HEC"});
        auto smallm = lemon::select::small_molecules(entry);
        // Pruning phase
        lemon::prune::identical_residues(entry, smallm);
        lemon::prune::cofactors(entry, smallm, lemon::common_cofactors);
        lemon::prune::cofactors(entry, smallm, lemon::common_fatty_acids);
        lemon::prune::keep_interactions(entry, smallm, hemegs, distance);
        // Output phase
        return pdbid + lemon::count::print_residue_names(entry, smallm);
    };
    auto collector = lemon::print_combine(std::cout);
    return lemon::launch(o, worker, collector);
}
```

**Listing S9:** (distance-based) C++ Lambda function to determine the number of small molecules which interact with a SAM molecule within a distance cutoff. This workflow is similar in spirit to the previous one. It was written by request of a user interested in the interaction of ligands with this cofactor.

```cpp
int main(int argc, char* argv[]) {
    lemon::Options o;
    auto distance = 6.0;
    o.add_option("--distance,-d", distance,
                 "Largest distance between a SAM group and a small molecule.");
    o.parse_command_line(argc, argv);
    auto worker = [distance](chemfiles::Frame entry,
                             const std::string& pdbid) {
        // Selection phase
        auto sam = lemon::select::specific_residues(entry, {"SAM"});
        auto smallm = lemon::select::small_molecules(entry);
        // Pruning phase
        lemon::prune::identical_residues(entry, smallm);
        lemon::prune::cofactors(entry, smallm, lemon::common_cofactors);
        lemon::prune::cofactors(entry, smallm, lemon::common_fatty_acids);
        lemon::prune::keep_interactions(entry, smallm, sam, distance);
        // Output phase
        return pdbid + lemon::count::print_residue_names(entry, smallm);
    };

    auto collector = lemon::print_combine(std::cout);
    return lemon::launch(o, worker, collector);
}
```

**Listing S10:** (distance-based) C++ Lambda function to find small molecules which do not interact with any water molecules within a distance cutoff. Water is an important consideration when predicting the pose of a ligand in a binding site and therefore many users may wish to find ligands which are within a given proximity to water.

```
int main(int argc, char* argv[]) {
    lemon::Options o;
    auto distance = 6.0;
    o.add_option("--distance,-d", distance,
                 "Largest distance between water and a small molecule.");
    o.parse_command_line(argc, argv);

    auto worker = [distance](chemfiles::Frame entry,
                             const std::string& pdbid) {
        // Selection phase
        auto waters = lemon::select::specific_residues(entry, {"HOH"});
        auto smallm = lemon::select::small_molecules(entry);
        // Pruning phase
        lemon::prune::identical_residues(entry, smallm);
        lemon::prune::cofactors(entry, smallm, lemon::common_cofactors);
        lemon::prune::cofactors(entry, smallm, lemon::common_fatty_acids);
        lemon::prune::remove_interactions(entry, smallm, waters, distance);
        // Output phase
        return pdbid + lemon::count::print_residue_names(entry, smallm);
    };
    auto collector = lemon::print_combine(std::cout);
    return lemon::launch(o, worker, collector);
}
```

**Listing S11:** (distance-based) C++ Lambda function to find small molecules which interact with an amino acid chemical group. These interactions are crucial to developing small-molecule therapeutics and are thus of great important to the medicinal chemistry community.

```
int main(int argc, char* argv[]) {
    lemon::Options o;
    auto distance = 6.0;
    o.add_option("--distance,-d", distance,
                 "Largest distance between a protein and a small molecule.");
    o.parse_command_line(argc, argv);
    auto worker = [distance](chemfiles::Frame entry,
                             const std::string& pdbid) {
        // Selection phase
        auto peptides = lemon::select::peptides(entry);
        auto smallm = lemon::select::small_molecules(entry);
        // Pruning phase
        lemon::prune::identical_residues(entry, smallm);
        lemon::prune::cofactors(entry, smallm, lemon::common_cofactors);
        lemon::prune::cofactors(entry, smallm, lemon::common_fatty_acids);
        lemon::prune::keep_interactions(entry, smallm, peptides, distance);
        // Output phase
        return pdbid + lemon::count::print_residue_names(entry, smallm);
    };

    auto collector = lemon::print_combine(std::cout);
    return lemon::launch(o, worker, collector);
}
```

**Listing S12:** (distance-based) C++ Lambda function to find small molecules which interact with a nucleic acid chemical group. This example was written by request from a user wishing to study the interactions between RNA and small-molecules.

```
int main(int argc, char* argv[]) {
    lemon::Options o;
    auto distance = 6.0;
    o.add_option("--distance,-d", distance,
                 "Largest distance between a nucleic-acid and a small
molecule.");
    o.parse_command_line(argc, argv);
    auto worker = [distance](chemfiles::Frame entry,
                             const std::string& pdbid) {
        // Selection phase
        auto nucleic_acids = lemon::select::nucleic_acids(entry);
        auto smallm = lemon::select::small_molecules(entry);
        // Pruning phase
        lemon::prune::identical_residues(entry, smallm);
        lemon::prune::cofactors(entry, smallm, lemon::common_cofactors);
        lemon::prune::cofactors(entry, smallm, lemon::common_fatty_acids);
        lemon::prune::keep_interactions(entry, smallm, nucleic_acids, distance);
        // Output phase
        return pdbid + lemon::count::print_residue_names(entry, smallm);
    };
    auto collector = lemon::print_combine(std::cout);
    return lemon::launch(o, worker, collector);
}
```

**Listing S13:** (complex) Lemon C++ Workflow to align all structures to a given reference structure using the TMalign algorithm and print the corresponding scores.

```
int main(int argc, char* argv[]) {

    lemon::Options o;

    auto reference = std::string("reference.pdb");

    o.add_option("--reference,-r", reference, "Protein or DNA to align to.")->

        check(CLI::ExistingFile);

    o.parse_command_line(argc, argv);

    chemfiles::Trajectory traj(reference);

    chemfiles::Frame native = traj.read();

    auto worker = [&native](chemfiles::Frame entry,

                            const std::string& pdbid) {

        std::vector<chemfiles::Vector3D> junk;

        auto tm = lemon::tmalign::TMscore(entry, native, junk);

        return pdbid + "\t" +

               std::to_string(tm.score) + "\t" +

               std::to_string(tm.rmsd) + "\t" +

               std::to_string(tm.aligned) + "\n";

    };

    auto collector = lemon::print_combine(std::cout);

    return lemon::launch(o, worker, collector);

}
```

**Listing S14:** (complex) Lemon C++ Workflow to calculate the docking score of all small-molecules with the surrounding environment using the scoring function published with AutoDOCK Vina.

```
int main(int argc, char* argv[]) {
    lemon::Options o(argc, argv);
    auto worker = [](chemfiles::Frame entry,
                     const std::string& pdbid) {
        // Selection phase
        std::list<size_t> smallm;
        if (lemon::select::small_molecules(entry, smallm) == 0) {
            return std::string("");
        }
        // Pruning phase
        lemon::prune::identical_residues(entry, smallm);
        lemon::prune::cofactors(entry, smallm, lemon::common_cofactors);
        lemon::prune::cofactors(entry, smallm, lemon::common_fatty_acids);
        // Output phase
        const auto& residues = entry.topology().residues();
        std::list<size_t> proteins;
        for (size_t i = 0; i < entry.topology().residues().size(); ++i) {
            proteins.push_back(i);
        }
        std::string result;
        for (auto smallm_id : smallm) {
            auto prot_copy = proteins;
            lemon::prune::keep_interactions(entry, smallm, prot_copy, 8.0);
            prot_copy.erase(std::remove(prot_copy.begin(), prot_copy.end(),
smallm_id));
            auto vscore =
                lemon::xscore::vina_score(entry, smallm_id, prot_copy);
            result += pdbid + "\t" +
```

```
                residues[smallm_id].name() + "\t" +
                std::to_string(vscore.g1) + "\t" +
                std::to_string(vscore.g2) + "\t" +
                std::to_string(vscore.hydrogen) + "\t" +
                std::to_string(vscore.hydrophobic) + "\t" +
                std::to_string(vscore.rep) + "\n";
        }
        return result;
    };
    auto collector = lemon::print_combine(std::cout);
    return lemon::launch(o, worker, collector);
}
```

**Listing S15:** (complex) C++ Lambda function to calculate all bond distances in the PDB.

```
// typedefs for binned data
typedef std::pair<std::string, size_t> BondStretchBin;
typedef std::map<BondStretchBin, size_t> StretchCounts;

using lemon::geometry::protein::bond_name;

int main(int argc, char* argv[]) {
    lemon::Options o;
    auto bin_size = 0.01;
    o.add_option("--bin_size,-b", bin_size, "Size of the length(stretch) bin.");
    o.parse_command_line(argc, argv);

    auto worker = [bin_size](chemfiles::Frame entry,
                             const std::string& pdbid) {
        StretchCounts bins;
        // Selection phase
        chemfiles::Frame protein_only;
        std::list<size_t> peptides;
        if (lemon::select::specific_residues(entry, peptides,
                                              lemon::common_peptides) == 0) {
            return bins;
        }
        lemon::separate::residues(entry, peptides, protein_only);
        const auto& bonds = protein_only.topology().bonds();
        for (const auto& bond : bonds) {
            std::string bondnm;
            try {
                bondnm = bond_name(protein_only, bond);
```

```
            } catch (const lemon::geometry::geometry_error& e) {
                auto msg = pdbid + ": " + e.what() + '\n';
                std::cerr << msg;
            }
            auto distance = protein_only.distance(bond[0], bond[1]);
            size_t bin = static_cast<size_t>(std::floor(distance / bin_size));
            BondStretchBin sbin = {bondnm, bin};
            auto bin_iterator = bins.find(sbin);
            if (bin_iterator == bins.end()) {
                bins[sbin] = 1;
                continue;
            }
            ++(bin_iterator->second);
        }
        return bins;
    };
    StretchCounts sc_total;
    auto collector = lemon::map_combine<StretchCounts>(sc_total);
    lemon::launch(o, worker, collector);
    for (const auto& i : sc_total) {
        std::cout << i.first.first << "\t"
                  << static_cast<double>(i.first.second) * bin_size << "\t"
                  << i.second << "\n";
    }
    return 0;
}
```

**Listing S16:** (complex) C++ Lambda function to calculate all bond angles in the PDB.

```
// typedefs for binned data
typedef std::pair<std::string, size_t> BondAngleBin;
typedef std::map<BondAngleBin, size_t> AngleCounts;

using lemon::geometry::protein::angle_name;

int main(int argc, char* argv[]) {
    lemon::Options o;
    auto bin_size = 0.01;
    o.add_option("--bin_size,-b", bin_size, "Size of the angle bin.");
    o.parse_command_line(argc, argv);
    auto worker = [bin_size](chemfiles::Frame entry,
                             const std::string& pdbid) {
        AngleCounts bins;
        // Selection phase
        chemfiles::Frame protein_only;
        std::list<size_t> peptides;
        if (lemon::select::specific_residues(entry, peptides,
                                             lemon::common_peptides) == 0) {
            return bins;
        }
        lemon::separate::residues(entry, peptides, protein_only);
        const auto& angles = protein_only.topology().angles();
        for (const auto& angle : angles) {
            std::string anglenm;
            try {
                anglenm = angle_name(protein_only, angle);
            } catch (const lemon::geometry::geometry_error& e) {
```

```
                auto msg = pdbid + ": " + e.what() + '\n';
                std::cerr << msg;
            }
            auto theta = protein_only.angle(angle[0], angle[1], angle[2]);
            size_t bin = static_cast<size_t>(std::floor(theta / bin_size));
            BondAngleBin sbin = {anglenm, bin};
            auto bin_iterator = bins.find(sbin);
            if (bin_iterator == bins.end()) {
                bins[sbin] = 1;
                continue;
            }
            ++(bin_iterator->second);
        }
        return bins;
    };
    AngleCounts sc_total;
    auto collector = lemon::map_combine<AngleCounts>(sc_total);
    lemon::launch(o, worker, collector);
    for (const auto& i : sc_total) {
        std::cout << i.first.first << "\t"
                  << static_cast<double>(i.first.second) * bin_size << "\t"
                  << i.second << "\n";
    }
    return 0;
}
```

**Listing S17:** (complex) C++ Lambda function to calculate all bond improper dihedrals in the PDB.

```
// typedefs for binned data
typedef std::pair<std::string, int> BondImproperBin;
typedef std::map<BondImproperBin, size_t> ImproperCounts;

using lemon::geometry::protein::improper_name;

int main(int argc, char* argv[]) {
    lemon::Options o;
    auto bin_size = 0.01;
    o.add_option("--bin_size,-b", bin_size, "Size of the improper-dihedral
bin.");
    o.parse_command_line(argc, argv);
    auto worker = [bin_size](chemfiles::Frame entry,
                             const std::string& pdbid) {
        ImproperCounts bins;
        // Selection phase
        chemfiles::Frame protein_only;
        std::list<size_t> peptides;
        if (lemon::select::specific_residues(entry, peptides,
                                             lemon::common_peptides) == 0) {
            return bins;
        }
        lemon::separate::residues(entry, peptides, protein_only);
        protein_only.set_cell(entry.cell());
        const auto& impropers = protein_only.topology().impropers();
        for (const auto& improper : impropers) {
            std::string impropernm;
            try {
```

```
                impropernm = improper_name(protein_only, improper);
            } catch (const lemon::geometry::geometry_error& e) {
                auto msg = pdbid + ": " + e.what() + '\n';
                std::cerr << msg;
            }
            auto theta = protein_only.out_of_plane(improper[0], improper[1],
                                                   improper[2], improper[3]);
            int bin = static_cast<int>(std::floor(theta / bin_size));
            BondImproperBin sbin = {impropernm, bin};
            auto bin_iterator = bins.find(sbin);
            if (bin_iterator == bins.end()) {
                bins[sbin] = 1;
                continue;
            }
            ++(bin_iterator->second);
        }
        return bins;
    };
    ImproperCounts sc_total;
    auto collector = lemon::map_combine<ImproperCounts>(sc_total);
    lemon::launch(o, worker, collector);
    for (const auto& i : sc_total) {
        std::cout << i.first.first << "\t"
                  << static_cast<double>(i.first.second) * bin_size << "\t"
                  << i.second << "\n";
    }
    return 0;
}
```

**Listing S18:** (complex) C++ Lambda function to calculate all bond dihedrals in the PDB.

```
// typedefs for binned data
typedef std::pair<std::string, int> BondDihedralBin;
typedef std::map<BondDihedralBin, size_t> DihedralCounts;

using lemon::geometry::protein::dihedral_name;

int main(int argc, char* argv[]) {
    lemon::Options o;
    auto bin_size = 0.01;
    o.add_option("--bin_size,-b", bin_size, "Size of the dihedral bin.");
    o.parse_command_line(argc, argv);
    auto worker = [bin_size](chemfiles::Frame entry,
                             const std::string& pdbid) {
        DihedralCounts bins;

        // Selection phase
        chemfiles::Frame protein_only;
        std::list<size_t> peptides;
        if (lemon::select::specific_residues(entry, peptides,
                                             lemon::common_peptides) == 0) {
            return bins;
        }
        lemon::separate::residues(entry, peptides, protein_only);
        protein_only.set_cell(entry.cell());
        const auto& dihedrals = protein_only.topology().dihedrals();
        for (const auto& dihedral : dihedrals) {
            std::string dihedralnm;
            try {
```

```
                dihedralnm = dihedral_name(protein_only, dihedral);
            } catch (const lemon::geometry::geometry_error& e) {
                auto msg = pdbid + ": " + e.what() + '\n';
                std::cerr << msg;
            }
            auto theta = protein_only.dihedral(dihedral[0], dihedral[1],
                                               dihedral[2], dihedral[3]);
            int bin = static_cast<int>(std::floor(theta / bin_size));
            BondDihedralBin sbin = {dihedralnm, bin};
            auto bin_iterator = bins.find(sbin);
            if (bin_iterator == bins.end()) {
                bins[sbin] = 1;
                continue;
            }
            ++(bin_iterator->second);
        }
        return bins;
    };
    DihedralCounts sc_total;
    auto collector = lemon::map_combine<DihedralCounts>(sc_total);
    lemon::launch(o, worker, collector);
    for (const auto& i : sc_total) {
        std::cout << i.first.first << "\t"
                  << static_cast<double>(i.first.second) * bin_size << "\t"
                  << i.second << "\n";
    }
    return 0;
}
```

**Listing S19:**(python) This workflow is a port of **Listing S6** and is an example of a 'simple' workflow that includes the 'selection' and 'pruning' of chemical groups. It illustrates how easy converting between Python and C++ implementations of lemon can be if one follows the recommend workflow development pipeline.

```
import lemon


class MyWorkflow(lemon.Workflow):
    def worker(self, entry, pdbid):
        import lemon
        smallm = lemon.select_small_molecules(entry, lemon.small_molecule_types,
10)


        # Pruning phase
        lemon.prune_identical_residues(entry, smallm)
        lemon.prune_cofactors(entry, smallm, lemon.common_cofactors)
        lemon.prune_cofactors(entry, smallm, lemon.common_fatty_acids)


        # Output phase
        return pdbid + lemon.count_print_residue_names(entry, smallm)
```

**Listing S20:**(python) This workflow is a Python port of **Listing S10** and again illustrates the similarites shared between the C++ and Python APIs.

```
import lemon


class MyWorkflow(lemon.Workflow):
    def worker(self, entry, pdbid):
        import lemon
        wat_name = lemon.ResidueNameSet()
        wat_name.append(lemon.ResidueName("HOH"))
        waters = lemon.select_specific_residues(entry, wat_name)
        smallm = lemon.select_small_molecules(entry, lemon.small_molecule_types,
10)

        # Pruning phase
        lemon.prune_identical_residues(entry, smallm)
        lemon.prune_cofactors(entry, smallm, lemon.common_cofactors)
        lemon.prune_cofactors(entry, smallm, lemon.common_fatty_acids)
        lemon.keep_interactions(entry, smallm, waters, 6.0)

        # Output phase
        return pdbid + lemon.count_print_residue_names(entry, smallm) + '\n'
```

**Listing S21:**(python) This workflow is a Python port of **Listing S17** and is an example of a 'complex' workflow implemented in Python. It is also an example of how to implement more functionality in the Python derived subclass.

```
from __future__ import print_function
import lemon


class MyWorkflow(lemon.Workflow):
    def __init__(self):
        import lemon
        lemon.Workflow.__init__(self) # This line is very important!
        self.dihedral_dict = {}


    def worker(self, entry, pdbid):
        import lemon
        import math


        protein_only = lemon.Frame()
        peptides = lemon.ResidueIDs()
        if (lemon.select_specific_residues(entry, peptides,
                                           lemon.common_peptides) == 0):
            return ""


        lemon.separate_residues(entry, peptides, protein_only)
        dihedrals = protein_only.topology().dihedrals()
        for dihedral in dihedrals:
            dihedralnm = ""
            try:
                dihedralnm = lemon.protein_dihedral_name(protein_only, dihedral,
lemon.proline_res)
            except lemon.GeometryError as error:
```

```python
                return pdbid + ": " + 'error' + '\n'
            theta = protein_only.dihedral(dihedral[0], dihedral[1],
                                          dihedral[2], dihedral[3])
            dbin = int(math.floor(theta / 0.01))
            sbin = (dihedralnm, dbin)
            if sbin in self.dihedral_dict:
                self.dihedral_dict[sbin] = self.dihedral_dict[sbin] + 1
            else:
                self.dihedral_dict[sbin] = 1
        return ""
    def finalize(self):
        for sbin, count in self.dihedral_dict.items():
            print(sbin[0], '\t', sbin[1], '\t', count)
```
